# How many cells resolve a perturbation direction? A closed-form, control-aware cell quota for single-cell perturbation screens

**DOI:** 10.64898/2026.10.02.754788

**Authors:** Tran Thai Huu Loc, Nguyen Thi Van

## Abstract

Single-cell perturbation screens increasingly infer mechanism from the direction of a treatment’s displacement in a shared transcriptomic embedding, yet no sample-size rule states how many cells that direction needs. We derive a closed-form, anisotropic cell quota for the treated cells required to hold the angular error of an estimated perturbation direction at a chosen tolerance. Because noise aligned with the effect changes only its length, the quota depends on the noise perpendicular to the effect, on the effect magnitude and on the control-pool size, and it diverges below a magnitude floor set by the control alone. The deterministic core is machine-checked in Lean 4. Across four public atlases (six screens) reprocessed through one embedding, the per-cell noise variance is near 1.0 on chemical and genome-wide genetic platforms. On Tahoe-100M, referenced to its plate-shared DMSO vehicle of about 3,100 cells, 55% of 56,827 conditions are control-pool-limited and only 11% are over-sampled at a 5.7-degree tolerance. Genome-wide CRISPRi knockdowns are instead detection- and depth-limited. A downsample-and-measure test on every atlas recovers the parameter-free angular-error slope. The calculator turns a pilot covariance, an effect magnitude and a control design into a per-condition cell budget and flags conditions that no treated depth can resolve.

## Introduction

Single-cell perturbation screens apply a genetic or chemical perturbation and read out the transcriptome of each treated cell, measuring how thousands of interventions alter gene expression at single-cell resolution [1–6]. Pooled CRISPR designs [7,8] and cell-resolved small-molecule screens [9,10] now assay perturbation libraries at a scale that is building reference atlases of cellular response [11,12]. A principal use of these atlases is to place an unannotated perturbation next to well-characterized ones and ask which it most resembles.

The comparison follows a standard workflow. Counts are normalized and projected into a shared low-dimensional embedding, typically a few dozen principal components of the log-normalized highly variable genes [13–16]. Each **condition** (one perturbagen at one dose in one cell line) is summarized by the centroid of its treated cells relative to matched controls, because per-cell tests conflate biological with technical replication [17–19]. The displacement between treated and control centroids is the perturbation’s effect. Mechanism is then read from the **direction** of that displacement and not from its size. Two treatments that move the transcriptome the same way are inferred to engage related programs, which is the single-cell analogue of connectivity-map reasoning [20,21].

This reliance on direction raises a question that has not been addressed directly. How many cells are needed to resolve a measured direction to a target angular precision? Two mechanistically distinct perturbations are separable only when each direction is resolved more tightly than the angle between them, so angular precision is the binding constraint and raw cell count is only a proxy for it. In practice the number is chosen heuristically. A conventional power calculation does not close the gap, because it assumes isotropic noise. Single-cell variance is instead concentrated on a few dominant axes reflecting cell cycle, lineage and technical variation [14,16], which is the spiked-covariance regime of high-dimensional principal-component analysis [22]. Whether a direction is easy or hard to orient depends on how it sits relative to those axes. Existing single-cell sample-size tools such as scPower [23], and the replicate-planning tradition inherited from bulk RNA-seq [24], address detection power and do not consider directional precision, and neither accounts for the control arm. Recent effort has gone into predicting [25–27] or harmonizing [11] perturbation responses, and no tool states how many cells a direction requires.

Here we derive a closed-form, anisotropic cell quota for direction estimation. It weights each embedding axis by how much it rotates the direction of interest, sets aside variance aligned with the signal, and carries the control-pool size explicitly, so that a small shared control emerges as its own binding constraint through a magnitude floor. We verify the rule symbolically, by simulation and, for its deterministic core, by a machine-checked proof. We then reprocess four public perturbation atlases, six screens across two modalities and three platforms, from raw counts through one fixed embedding, map the per-condition sufficiency spectrum of each, and test the underlying angular-error law by downsampling every atlas on its own cells. Across platforms and modalities the per-cell noise variance transfers in a fixed embedding, so a pilot on one platform calibrates the rule for another. The calculator and all derived fixtures are released as open-source software.

## Methods

### The angular-error model and the cell quota

Work in a fixed *d*-dimensional embedding. Treated cells 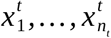 and control cells 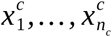 are independent draws with means *μ*_*t*_, *μ*_*c*_ and per-cell covariances *Σ*_*t*_, *Σ*_*c*_. The estimands are the perturbation vector *v*=*μ*_*t*_ *−μ*_*c*_, its magnitude *m*=‖*v*‖ and its unit direction *u*=*v* / *m*. The estimator subtracts the two centroids, 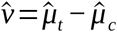 with 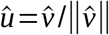, so by the multivariate central limit theorem 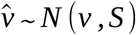 with the two-arm sampling covariance

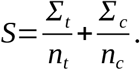

The angular error is *θ*=arccos(*û*^*⊤*^ *u*). Sampling error 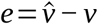 splits into a component along *u*, which changes only the length of ^*v*, and a perpendicular component, which alone rotates it (Figure 2). With the tangent-space projector *P*=*I −uu*^*⊤*^, the Jacobian of the normalization map at *v* is exactly *P*/ *m*, and a first-order Delta expansion in the high signal-to-noise regime *ρ*^2^ *≡m*^2^ /(*u*^*⊤*^ *Su*) *≫* 1 [28– 31] gives

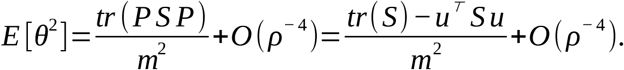

From the total sampling noise one subtracts the part aligned with the signal, and only the residual perpendicular noise contributes. In a PCA basis with per-component within-condition variances *ℓ*_*k*_,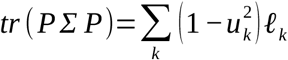, which reduces to for isotropic noise. The relative error of the first-order rule is 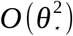, below 1% at *θ* _*\**_ =0.1 rad (Supplemental Note S1).

Setting 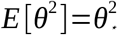 with a common *Σ* and solving for the treated arm gives the **two-arm cell quota**

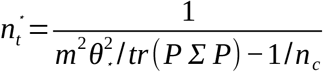

which reduces to the matched equal-arm value 2 *tr*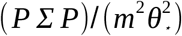 when *n*_*c*_=*n*_*t*_ and to the large-control limit *tr* 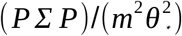 as *n*_*c*_ *→∞*. The denominator vanishes at the **control-pool floor**

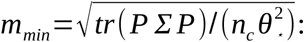

below it the control’s own sampling noise rotates the direction by more than 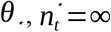, and the only remedies are a larger control pool or a looser tolerance. A Laurent–Massart tail bound extends the mean quota to a confidence quota 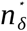 with *Pr* (*θ*>*θ*_*⋆*_)*≤δ* (Supplemental Note S2). When a centroid is instead sub-sampled from *N*_0_ cells already acquired, the sampling covariance carries the finite-population factor (1/ *n−*1 / *N* _0_) [32].

### Datasets and preprocessing

Every dataset was reprocessed from raw counts with one recipe: normalize each cell to 10^4^ counts, log (1+ *x*), 2,000 highly variable genes, PCA to *d*=50 [13]. The PCA basis is fitted on pooled single cells. The per-cell covariance *Σ* is then estimated from individual embedded cells as the pooled within-condition residual covariance (each condition centred on its own mean), and only afterwards is each condition summarized by the coordinate mean of its cells. Aggregating counts before PCA would collapse *Σ* to zero and remove the functional the quota depends on. The chemical pipelines store per-condition sufficient statistics and therefore carry a diagonal *Σ* (per-component variances). The genetic pipelines carry the full 50 *×*50 matrix.

**Tahoe-100M** [12] comprises 100,648,790 cells from 379 small molecules at three doses across 50 cancer cell lines on 14 96-well plates, each well pooling all lines under one treatment and computationally demultiplexed. Of these, 95,624,334 cells pass quality control, giving 56,827 (drug *×* dose *×* line) conditions with a median of *N*_0_=1, 296 cells. Each plate carries two or three DMSO_TF vehicle wells, which are pooled to form one vehicle centroid per (plate *×* line). **EmeraldBay** [33,34] is a second screen on the same Vevo Mosaic platform read at five days instead of 24 hours (1,831,648 cells, 52 lines, 91 treatments plus two vehicle samples), giving 4,912 non-empty (sample *×* line) conditions with a median depth of 240 cells. Its highly variable gene set is frozen for embedding stability. **X-Atlas/Orion** [35] is a genome-wide dual-sgRNA CRISPRi Perturb-seq atlas on the Xaira FiCS platform (HCT116 and HEK293T, 18,903 target genes, about 8 million cells) with a large pooled non-targeting (NTC) control. Its condition is a (gene *×* line) knockdown. The **TRADE** essential-gene screens [36] (GEO GSE264667) in Jurkat and HepG2 cover 2,393 common-essential genes with about 2.5 *×*10^5^ cells per line and a median of 48–85 cells per gene, using the same design on a third platform. Dataset sizes, controls and calibration constants are collected in Supplemental Table S1.

### Units, controls and regime classification

All six screens are computed by one engine under identical conventions, and nothing branches on dataset or modality. The counting unit is the condition defined above. The spectrum is computed over all conditions with no cell-count filter, except that a genetic knockdown needs at least 25 cells to have an estimable centroid. Each condition is referenced to its **real** control pool of size *n*_*c*_: the plate-shared, pooled DMSO_TF vehicle for Tahoe-100M (median *n*_*c*_ *≈* 3,113 cells, shared by a median of 94 conditions), the per-line mean for EmeraldBay (*n*_*c*_ *≈* 2.3 *×*10^4^), and the pooled NTC centroid for the genetic screens (1.7 *×*10^5^ to 2.2 *×*10^5^ on Orion, 4.4 *×*10^3^ to 1.2 *×*10^4^ on TRADE). Referencing Tahoe-100M to its per-line mean instead, a different estimand, is reported as a sensitivity analysis.

For every condition the squared magnitude is bias-corrected for the finite-sample floor, 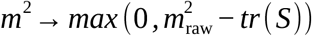 with *tr* (*S*)=*tr* (*Σ*)(1/ *n*_*t*_+1/ *n*_*c*_). A condition is **detectable** when 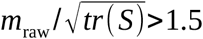. The quota is the two-arm form above at *θ*_*⋆*_ =0.1 rad (5.73°). Each condition is assigned to one of five regimes: OVER (*n*^*⋆*^ < *N*, already resolvable), UNDER (treated-depth-limited), GHOST (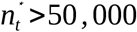, a feasibility convention), POOL-LIMITED 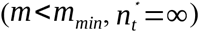 or NOT-DETECTABLE. For the genetic screens the over/under split is also reported among detectable knockdowns, because for the rest *m* is not measurable.

### Downsample-and-measure protocol

The angular-error law is tested on each atlas’s own cells. For each group with *N ≥* 400 acquired cells the reference direction is that of the full-depth centroid relative to the per-line mean. For each depth *n* on a geometric grid capped at *N* / 2, *R* subsamples are drawn without replacement, each yielding a direction *û* _*n*_ and an angle *θ*(*n*)=arccos 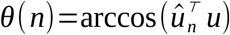. Because each subsample is drawn from the same *N* cells that define the reference, the prediction is *θ*^2^ (*n*)=*tr* (*P Σ P*)/ *m*^2^ *⋅* (1/ *n−*1/ *N*), so regressing the realized *θ*^2^ on (1/ *n−*1 / *N*) through the origin tests linearity with a slope equal to the parameter-free *tr* (*PΣ P*)/ *m*^2^. Groups are stratified by the theory’s validity criterion *ρ*^2^=*m*^2^ /(*u*^*⊤*^ *Σ u*) *≥* 3, and the predicted low signal-to-noise breakdown is quantified by the Spearman correlation between the slope deficit and 1/ *ρ*^2^. A separate decision test downsamples every condition predicted OVER to its quota and checks that the realized root-mean-square angle stays within 1.05 *θ*_*⋆*_. This threshold of 400 cells applies to the held-out tests only, never to the spectrum.

### Verification and software

The quota is verified four ways. A SymPy symbolic Jacobian of the normalization map matches *P*/ *m* exactly (residual 2.8 *×*10^*−* 17^). Monte-Carlo simulation (*K*=50,000) reproduces *E* [*θ*^2^] to 0.13% and the second-order correction to below 0.5%. The tail bound’s empirical exceedance is 0.34% at *δ* =10 %. The deterministic core (the Jacobian, the trace identity *tr* (*PΣ P*)=*tr Σ −u*^*⊤*^ *Σ u* and its isotropic and diagonal forms) is machine-checked in Lean 4/Mathlib with no unproven step. The probabilistic results are not formalized. Analyses are estimation-based and no significance thresholds are applied. The embedding was built with scanpy 1.12.2 on Python 3.13 (numpy 2.4.4, scipy 1.18.0, scikit-learn 1.9.0, anndata 0.12.18). The calculator, a 26-test suite and the reference checks run in continuous integration on Python 3.10–3.12 (Supplemental Methods).

## Results

### Overview of the framework

Figure 1 summarizes the study. A single closed-form quota is derived from the geometry of the angular error, verified independently of any dataset, and carried by one from-raw recompute to four perturbation atlases (six screens, two modalities). For each condition the calculator returns the required treated depth, or reports that no treated depth suffices because the control pool is too small. A tolerance sweep (Table 2) shows how many conditions each looser target would free.

**Figure 1.**
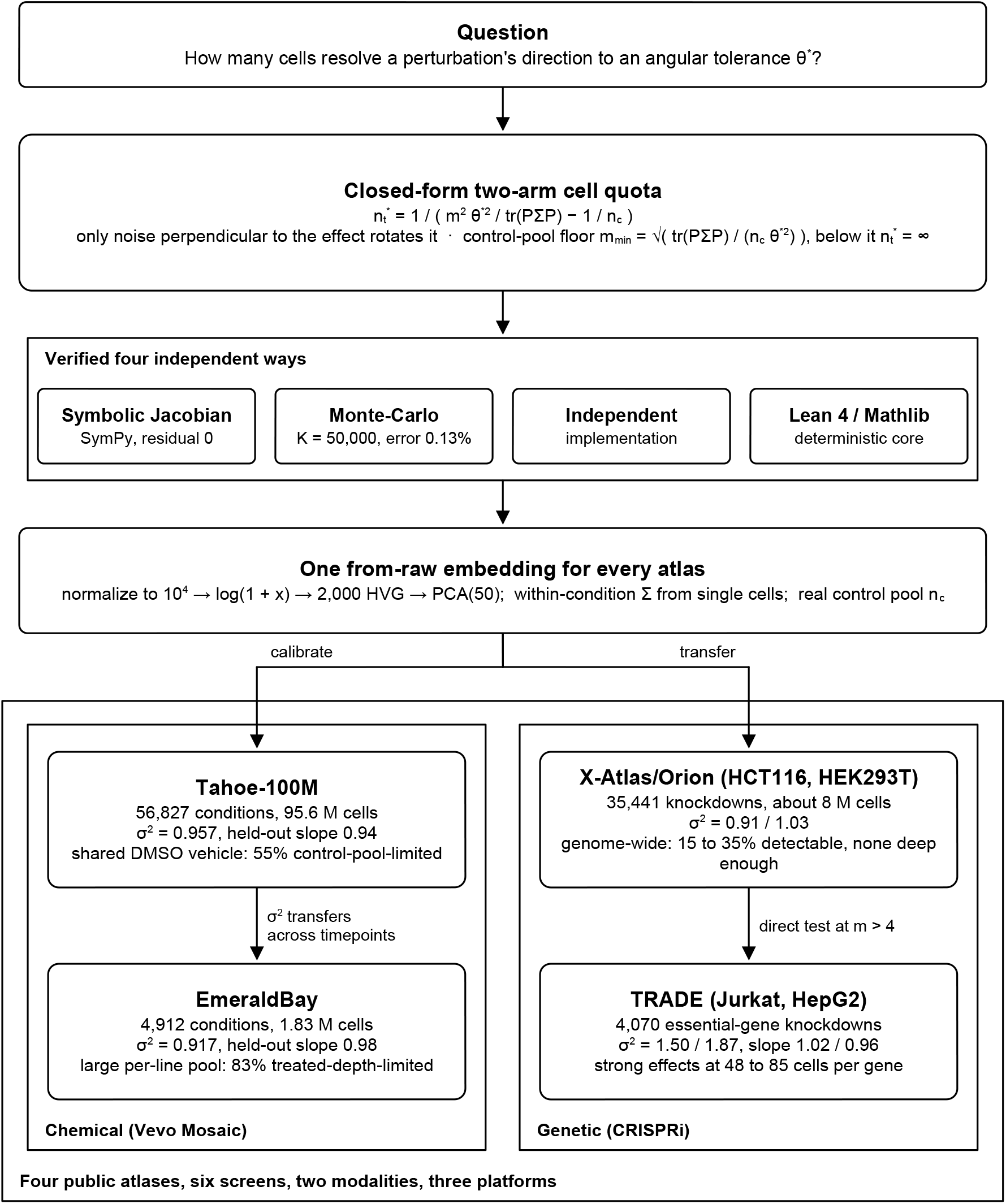
Study overview. One closed-form two-arm cell quota, 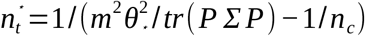, is derived from the geometry of the angular error, verified four independent ways, and carried by one from-raw embedding to four perturbation atlases (six screens). The quota is calibrated on Tahoe-100M and applied to EmeraldBay on the chemical side, and transferred to X-Atlas/Orion and tested on the TRADE screens on the genetic side. Each dataset box reports its noise variance and headline result.

**Figure 2.**
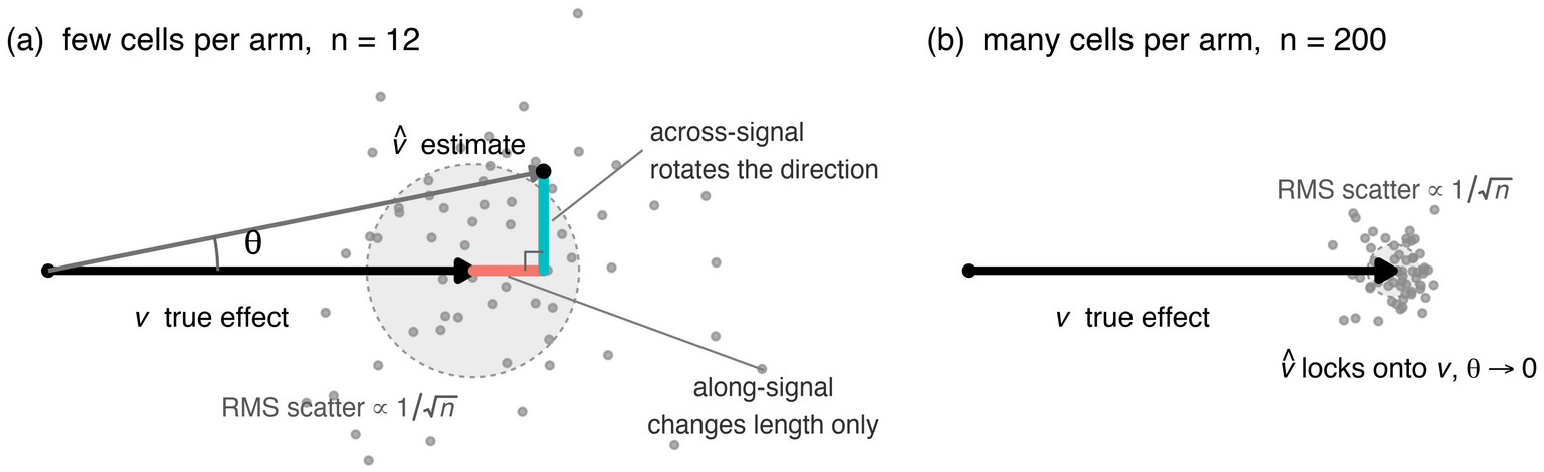
Geometry of the angular error. (a) With few cells per arm, the true effect vector *v* and a noisy estimate 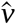 differ by a sampling error with an along-signal part, which changes only the length, and a perpendicular part, which rotates the direction by *θ*. The shaded disc is the root-mean-square radius of the cloud of estimates. (b) With many cells the cloud, and hence the angle, shrinks as 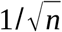. The quota controls only the perpendicular component.

### Only perpendicular noise rotates a direction

The estimand is the perturbation direction *u*, estimated by the unit vector of the centroid difference. Moving 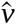 along its own direction changes its length but not its orientation, so only the transverse component of sampling noise contributes to the angle (Figure 2). Because *tr* (*PΣ P*) keeps only transverse noise, the effective noise dimension *d*_eff_ =(*tr PΣ P*)^2^/ *tr* ((*P Σ P*)^2^) is well below *d −*1 (12.2 in the Monte-Carlo check), and the quota depends on the data through this quantity instead of on the ambient dimension. With the Tahoe-100M calibration (*d*=50, within-condition *σ* ^2^=0.9567, *θ* =0.1 rad) the large-control limit becomes 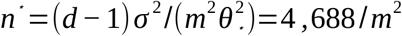 . A strong cytotoxic with *m*=15 then needs about 20 cells, a median condition about 3,900, and halving the magnitude quadruples the requirement.

### The sufficiency spectrum of Tahoe-100M is set by its control pool

Applying the quota to all 56,827 (drug *×* dose *×* line) conditions of Tahoe-100M, each referenced to its matched vehicle, yields the spectrum in Figure 3 and Table 1. The dominant regime is control-pool limitation. Of all conditions, 55.4% are pool-limited 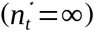, 5.6% are not detectable, 25.2% are treated-depth-limited, 2.4% are ghosts, and only 11.4% are over-sampled. Among conditions with a finite quota the median is 3,622 cells against a median acquired depth of 1,296. One pooled DMSO_TF centroid of about 3,113 cells per (plate *×* line) serves a median of 94 conditions, and its own sampling noise sets a median floor *m*_*min*_=1.22 that lies above the median bias-corrected effect magnitude of 1.096 (raw 1.14). For most conditions the control alone rotates the direction by more than *θ*_*⋆*_, so no treated depth resolves them. The conclusion is unchanged under the coarser marginal variance (Supplemental Note S3).

**Table 1.** Sufficiency spectrum across six screens (*θ*_*⋆*_ =0.1 rad, one engine). Percentages are over all conditions and sum to 100 with not-detectable. *σ* ^2^ is the within-condition per-cell variance, C the large-control bound 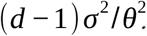, med. *m* the bias-corrected magnitude and *n*_*c*_ the real control pool (median for Tahoe).

| screen | unit | conditions | $n_c$ | $\sigma^2$ | $C$ | det. % | over % | under % | ghost % | pool-lim % | not-det % | med. $m$ | med. depth |
| --- | --- | --- | --- | --- | --- | --- | --- | --- | --- | --- | --- | --- | --- |
| Tahoe-100M | drug × dose × line | 56,827 | 3,113 | 0.9567 | 4,688 | 94.4 | 11.4 | 25.2 | 2.4 | 55.4 | 5.6 | 1.096 | 1,296 |
| EmeraldBay | drug × dose × line | 4,912 | 22,805 | 0.9174 | 4,495 | 87.6 | 2.7 | 82.5 | 2.0 | 0.5 | 12.4 | 1.175 | 240 |
| Orion HCT116 | gene × line | 17,585 | 165,562 | 0.9133 | 4,475 | 14.7 | 0.0 | 14.7 | 0.0 | 0.0 | 85.3 | 0.141 | 155 |
| Orion HEK293T | gene × line | 17,856 | 218,838 | 1.0334 | 5,063 | 35.3 | 0.0 | 35.3 | 0.0 | 0.0 | 64.7 | 0.351 | 204 |
| TRADE Jurkat | gene × line | 2,187 | 11,514 | 1.4975 | 7,338 | 70.6 | 0.8 | 68.3 | 0.8 | 0.7 | 29.4 | 1.841 | 85 |
| TRADE HepG2 | gene × line | 1,883 | 4,380 | 1.8667 | 9,147 | 60.9 | 2.7 | 56.7 | 0.3 | 1.2 | 39.1 | 2.249 | 48 |

**Table 2.** The control-pool limit versus angular tolerance (Tahoe-100M, real shared DMSO vehicle). As *θ*_*⋆*_ loosens the floor *m*_*min*_ falls as 1/ *θ*_*⋆*_, the pool-limited fraction collapses and the resolvable fraction grows.

| $\theta$ (rad) | $m_{min}$<br>(median) | pool-limited % | over % | under % | ghost % | not-detectable % |
| --- | --- | --- | --- | --- | --- | --- |
| 0.10 ($5.7^\circ$) | 1.22 | 55.4 | 11.4 | 25.2 | 2.4 | 5.6 |
| 0.15 ($8.6^\circ$) | 0.82 | 31.5 | 27.9 | 33.3 | 1.7 | 5.6 |
| 0.20 ($11.5^\circ$) | 0.61 | 17.9 | 44.8 | 30.7 | 0.9 | 5.6 |
| 0.25 ($14.3^\circ$) | 0.49 | 11.1 | 58.4 | 24.4 | 0.5 | 5.6 |
| 0.30 ($17.2^\circ$) | 0.41 | 7.2 | 68.1 | 18.8 | 0.2 | 5.6 |

**Figure 3.**
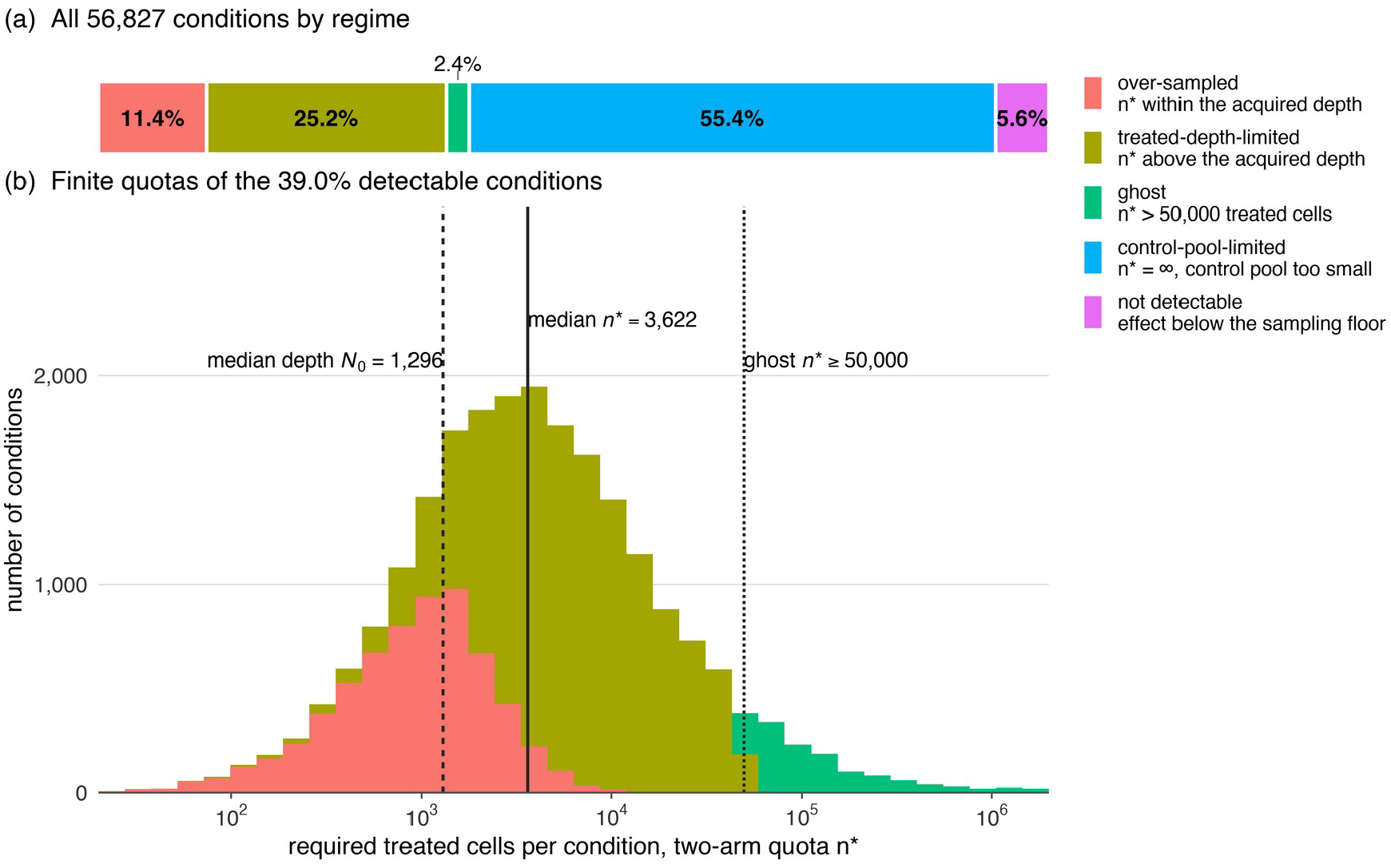
Sufficiency spectrum of Tahoe-100M against its pooled shared DMSO vehicle (*θ*_*⋆*_ =0.1 rad). (a) All 56,827 conditions by regime. (b) Finite quotas *n*^*⋆*^ of the 39.0% detectable conditions, stacked by regime with the colours of (a), on a logarithmic axis truncated at 2 *×*10^6^. Vertical lines mark the median acquired depth *N*_0_=1, 296, the median finite quota *n*^*⋆*^ =3, 622 and the ghost threshold *n*^*⋆*^ =50, 000. The control-pool-limited majority has no finite quota and appears only in (a).

Resolved by compound (Supplemental Table S2), 240 of 379 drugs have a median magnitude below the floor and are pool-limited in more than half of their 150 conditions. Only 9 drugs are over-sampled in a majority of their conditions, and 23 in none. The over-sampled drugs are the strongest displacers, the protein-synthesis inhibitors harringtonine and homoharringtonine, the proteasome inhibitor bortezomib and the HDAC inhibitor panobinostat (median *m ≈* 3.4–5.6, *n*^*⋆*^ *≈* 125–340), whereas translation-initiation and receptor-tyrosine-kinase inhibitors such as 4EGI-1 and (S)-crizotinib (median *m ≤*1.3) are under-sampled or pool-limited almost everywhere. Dose and cell line modulate the regime. Homoharringtonine in NCI-H460 moves from *m*=9.6 (*n*^*⋆*^ =51) at 0.05 µM to *m*=14.9 (*n*^*⋆*^ =21) at 5 µM, and across the panel the over-sampled fraction rises from 6.9% at the lowest dose to about 13.5% at the two higher doses (Supplemental Table S3). The same drug at 5 µM reaches only *m*=3.0 in NCI-H661, whose plate vehicle holds about 210 cells (*m*_*min*_=4.7), so it is pool-limited there. Because magnitude is set jointly by compound and context, quotas must be calibrated per condition from the measured magnitude and cannot be inferred from mechanism labels.

The choice of reference also matters. Referencing each condition to its per-line mean instead, the average over that line’s roughly 94 treatments, removes the floor (*m*_*min*_ *≈* 0.05) and gives 21.7% over-sampled with no pool limitation. That reference measures “difference from the average drug” and not “difference from untreated”, and it claims a control no real experiment has, so we report it only as a sensitivity analysis. The vehicle is the pool the direction estimand actually names. Directly downsampling the 6,467 conditions predicted over-sampled against the vehicle to their quota confirms the classification, with 6,442 (99.6%) reaching a realized root-mean-square angle within 1.05 *θ*_*⋆*_. The 25 exceptions (realized *≤* 6.5° against a 5.7° target) are strong-effect conditions, 23 of them in the single most responsive line, NCI-H460, which points to a within-condition covariance for strong responders that departs from the atlas-wide *Σ*. Because subsample and reference share the control centroid, this test exercises only the treated-arm term of the quota.

### Tolerance and control-pool size both act on the floor

Because 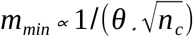, a condition below the floor can be freed either by enlarging the shared control or by accepting a coarser direction. Sweeping the tolerance on Tahoe’s vehicle (Table 2), loosening from 5.7° to 17° drops the pool-limited fraction from 55% to 7% and raises the resolved (over-sampled) fraction from 11% to 68%, with no extra cells. The not-detectable 5.6% is unmoved, because detectability is a signal-to-noise property independent of the angular target. The control-pool limit can therefore be designed around. A shared vehicle or non-targeting control must scale its pool with the number of conditions it references, or the screen buys treated precision it cannot realize. On this atlas, enlarging the shared DMSO pool would resolve more conditions than enlarging any treated arm.

### Consequences for screen design

#### Compression

The quota marks the regime in which centroid summaries stay stable under downsampling. On EmeraldBay, reducing an over-sampled group (DMSO_T0 *×* HS-578T, *N*_0_=1, 067, *n*^*⋆*^ =92) to its quota with 300 random replicates changed its centroid–baseline distance by *≤* 0.6 % and its cosine similarity to its three nearest neighbours by *≤* 0.005. The identical 92-cell reduction of an under-sampled group (gemcitabine *×* MIA PaCa-2, *N*_0_=280, *n*^*⋆*^ =42, 666) increased the distance by 19% and dropped the cosines by 0.10–0.12, with a tenfold larger replicate spread (Supplemental Table S4). Which conditions may be compressed is known in advance from *m* and *Σ*. In the wet lab the surplus of an over-acquired condition can be spent on multiplexing.

Homoharringtonine at 5 µM in NCI-H460 (*N*_0_=6, 060, *m*=14.9, *n*^*⋆*^ =21) carries a roughly 290-fold gain.

#### Replication

Replicates are largely absent in Tahoe-100M (86.3% of drug–dose combinations occupy one plate), and even among conditions with a finite quota the shortfall between quota and acquired depth is about 2.8-fold. The pool-limited majority cannot be closed by treated cells at all.

#### Budget and reliability

Where the control is large, the treated allocation is what can be optimized. For a 600-condition screen (100 drugs *×* 3 doses *×* 2 lines) drawing magnitudes from the Tahoe distribution under the per-line-mean reference, flat loading at the 90th-percentile quota (10,973 cells per condition) needs 6.6 M treated cells. Quota-guided allocation capped at 10,000 needs 2.4 M, a 2.8-fold reduction (Table 3). Against the real shared vehicle, however, treated allocation reaches only the 37% of conditions that a finite treated depth can resolve, and in this pool-limited regime magnitude-adaptive and budget-matched flat allocation recover the drug-similarity graph indistinguishably (*k* -nearest-neighbour Jaccard within 0.006 at every budget). A budget-matched test of mechanism-of-action label recovery on that graph did not exceed the majority-class baseline even at full depth (accuracy 0.062 against 0.142). On this atlas the quota is useful for deciding which conditions can be trusted and not for reallocating cells. A direction is resolved only when it clears the vehicle floor and *N*_0_ *≥n*^*⋆*^. This holds for 11.4% of conditions at *θ*_*⋆*_ =0.1 rad, 44.8% at 0.2 rad and 68.1% at 0.3 rad, and only 7.5% of *k* =10 neighbour edges at 0.1 rad connect two resolved directions (38.6% and 64.0% at 0.2 and 0.3 rad). At a stringent tolerance most pairwise similarity calls rest on at least one under-resolved direction, and the calculator flags those edges in advance.

**Table 3.**
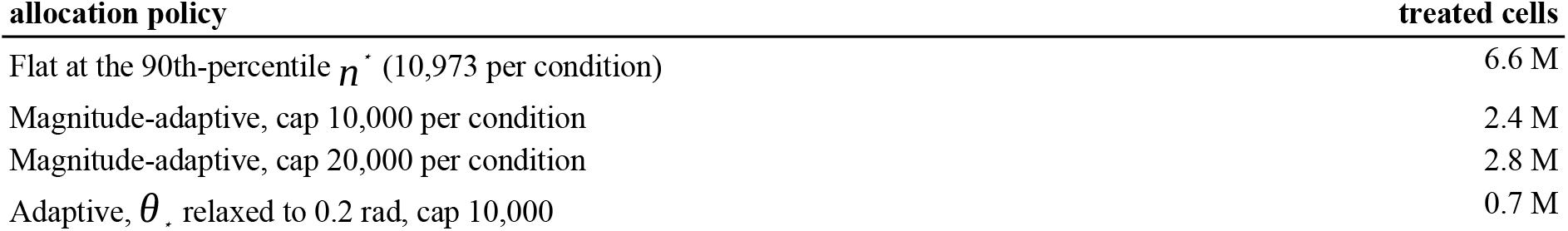
Screen-budget comparison for a 600-condition screen under the per-line-mean reference. The saving is the ratio of an adaptive row to the flat 90th-percentile row.

### Genetic perturbation: detection-limited genome-wide, depth-limited for essential genes

The quota applies unchanged to genetic perturbation because it is a statement about centroid-direction estimation. On X-Atlas/Orion the within-condition variance is *σ* ^2^=0.913 (HCT116) and 1.033 (HEK293T), in the same band as the chemical atlases (0.957 and 0.917) despite a different perturbation type, platform and laboratory. What transfers is the measurement noise. With a large pooled NTC control the quota approaches its large-control bound (4,475 and 5,063 per *m*^2^), so the operating constants of all screens referenced to a large pool fall in one 4,500–9,100 band (Table 1). The genome-wide result has two parts (Table 4). Only 14.7% (HCT116) and 35.3% (HEK293T) of knockdowns clear the detection floor, because a genome-wide library is dominated by weak non-essential knockdowns (median bias-corrected *m* 0.14 and 0.35) acquired at only 155–204 cells. Among detectable knockdowns almost none is deep enough. Only 0.0% and 0.1% are over-sampled, at a median deficit of 35- and 17-fold. The strongest knockdowns are core essential machinery: ribosomal proteins (RPL31, RPL18, RPL38, RPS20), ribosome biogenesis factors (MAK16, UTP15, NOL8, HEATR1), RNA polymerase II subunits (POLR2C, POLR2I), splicing factors (SF3B5, CPSF4), NUP93 and EIF2S3.

**Table 4.**
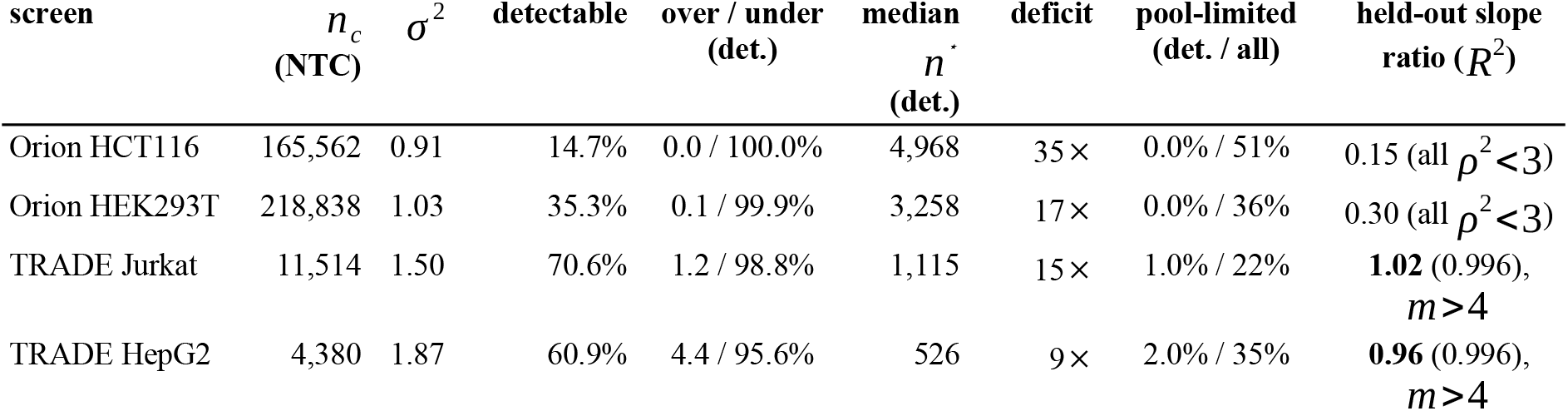
Genetic-perturbation sufficiency spectrum (*θ*_*⋆*_ =0.1 rad). *Detectable* is the fraction of knockdowns clearing the sampling floor (signal-to-noise > 1.5). Over/under, median *n*^*⋆*^ and deficit are among detectable knockdowns. Pool-limited is given among detectable and among all knockdowns. The last column is the held-out slope ratio (Table 5), recovered by the strong essentialgene knockdowns (*m* >4).

**Table 5.** Held-out downsample-and-measure test, one row per dataset, all groups with *N ≥* 400 cells. *Median ratio (all)* is low because most eligible groups are weak, whereas *median ratio (ρ*^2^ *≥*3) restricts to the first-order regime. The last column is the rank correlation between the slope deficit and 1/ *ρ*^2^. For the TRADE screens the slope is recovered at *m* >4 (Table 4).

| dataset | groups ($N \geq 400$) | median ratio (all) | $n(\rho^2 \geq 3)$ | median ratio ($\rho^2 \geq 3$) | median $R^2$ | Spearman deficit vs $1/\rho^2$ |
| --- | --- | --- | --- | --- | --- | --- |
| Tahoe-100M | 56,195 | 0.68 | 1,790 | 0.94 | 0.999 | 0.57 |
| EmeraldBay | 958 | 0.41 | 25 | 0.98 | 0.999 | 0.81 |
| Orion HCT116 | 1,059 | 0.15 | 0 | n/a (max $\rho^2=1.3$) | n/a | 0.02 |
| Orion HEK293T | 2,670 | 0.30 | 0 | n/a (max $\rho^2=2.0$) | n/a | 0.12 |
| TRADE Jurkat | 38 | 0.50 | 35 | 0.55 | 0.932 | 0.87 |
| TRADE HepG2 | 9 | 0.80 | 8 | 0.82 | 0.990 | 0.95 |

The TRADE essential-gene screens occupy the opposite regime. Their knockdowns are strong (median *m* 1.84 and 2.25, above Tahoe’s drugs) and mostly detectable (70.6% Jurkat, 60.9% HepG2), but they were sequenced shallowly (median 48–85 cells per gene), so again almost none is deep enough (1.2% and 4.4% of detectable knockdowns over-sampled). Their *σ* ^2^ is higher (1.50 and 1.87) because strong and diverse responses spread variance across many components, and their NTC pools are small (11,514 and 4,380 cells), so the pool floor is appreciable (HepG2 *m*_*min*_=1.39) even though it binds on only 1.0–2.0% of detectable knockdowns. The magnitude distributions and regime splits of all datasets are compared in Figure 4.

**Figure 4.**
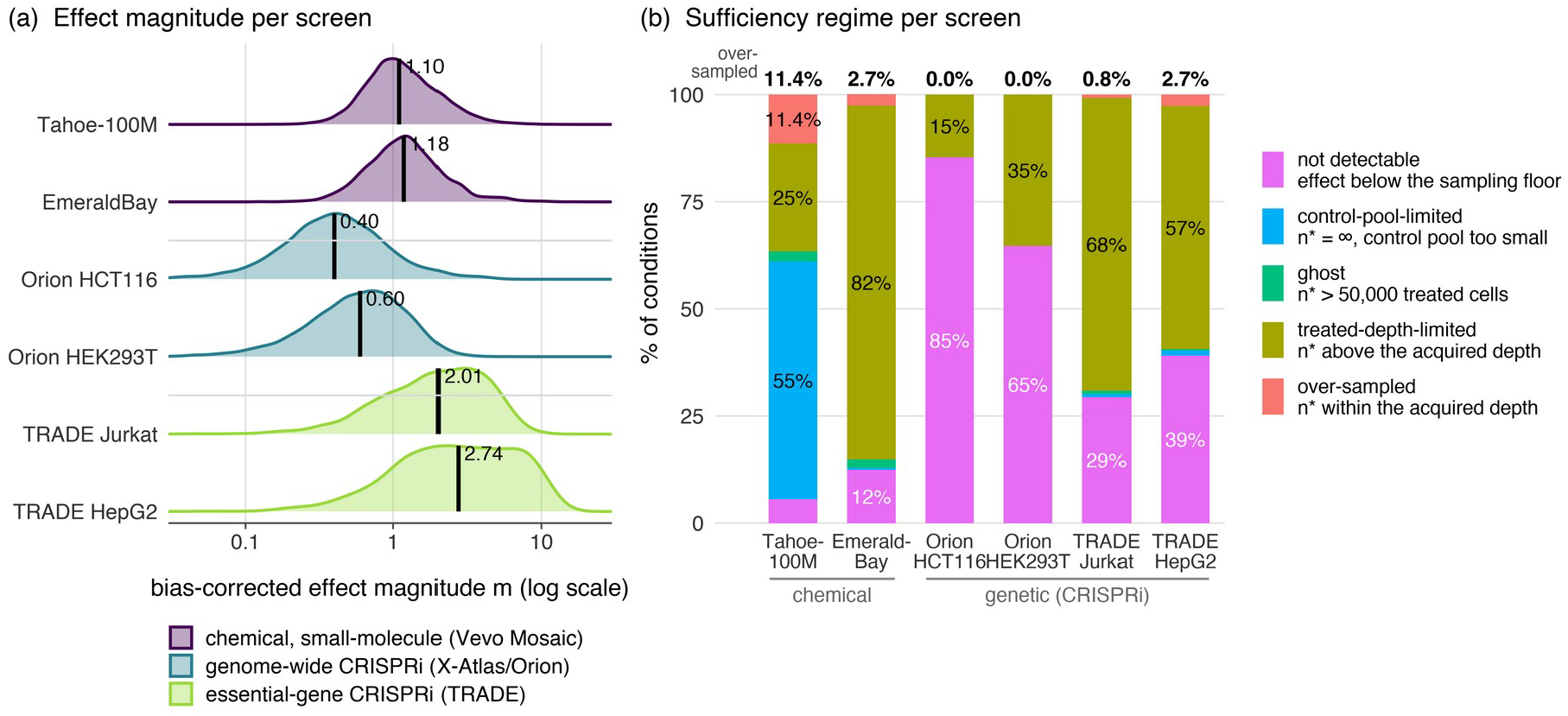
Cross-modality summary at *θ*_*⋆*_ =0.1 rad. (a) Distribution of the bias-corrected effect magnitude *m*, one row per screen, coloured by modality, with the median marked. Genome-wide knockdowns are several-fold weaker than drugs and essential-gene knockdowns stronger. (b) Sufficiency regime per screen as fractions of all conditions, with the regime colours of Figure 3 and the over-sampled fraction printed above each bar. The numbers are those of Table 1. EmeraldBay magnitudes are referenced to the per-line mean and sit on a different baseline.

### The angular-error law holds on every atlas where the signal is strong enough

The downsample-and-measure protocol tests the law itself, separately on every group of each atlas with at least 400 cells (Figure 5, Table 5). Wherever a dataset contains groups in the first-order regime (*ρ*^2^ *≥*3), the parameter-free slope *tr* (*PΣ P*)/ *m*^2^ is recovered at a median realized/predicted ratio near one with median *R*^2^ *≈* 0.999. The ratio is 0.94 on Tahoe-100M (1,790 of 56,195 groups) and 0.98 on EmeraldBay (25 of 958). Below that regime the deficit grows monotonically with 1/ *ρ*^2^ on every dataset that spans a signal-to-noise range (Spearman 0.57–0.95). The genome-wide Orion lines contain no group above *ρ*^2^=3 at all (maximum 1.3 and 2.0), so their tests land wholly in the breakdown regime (median ratio 0.15 and 0.30), which reflects the genome-wide magnitude distribution and not the law. The essential-gene TRADE screens reach the strong regime, where restricting to the knockdowns the theory addresses (*m* >4) recovers the slope directly in the genetic modality (median ratio 1.02 Jurkat, 0.96 HepG2, *R*^2^=0.996). The decision the quota makes also holds. All 706 EmeraldBay groups predicted over-sampled at *θ*_*⋆*_ =0.20 rad, and all 13,964 Tahoe conditions predicted over-sampled under the per-line-mean reference, meet the tolerance when downsampled to *n*^*⋆*^ .

**Figure 5.**
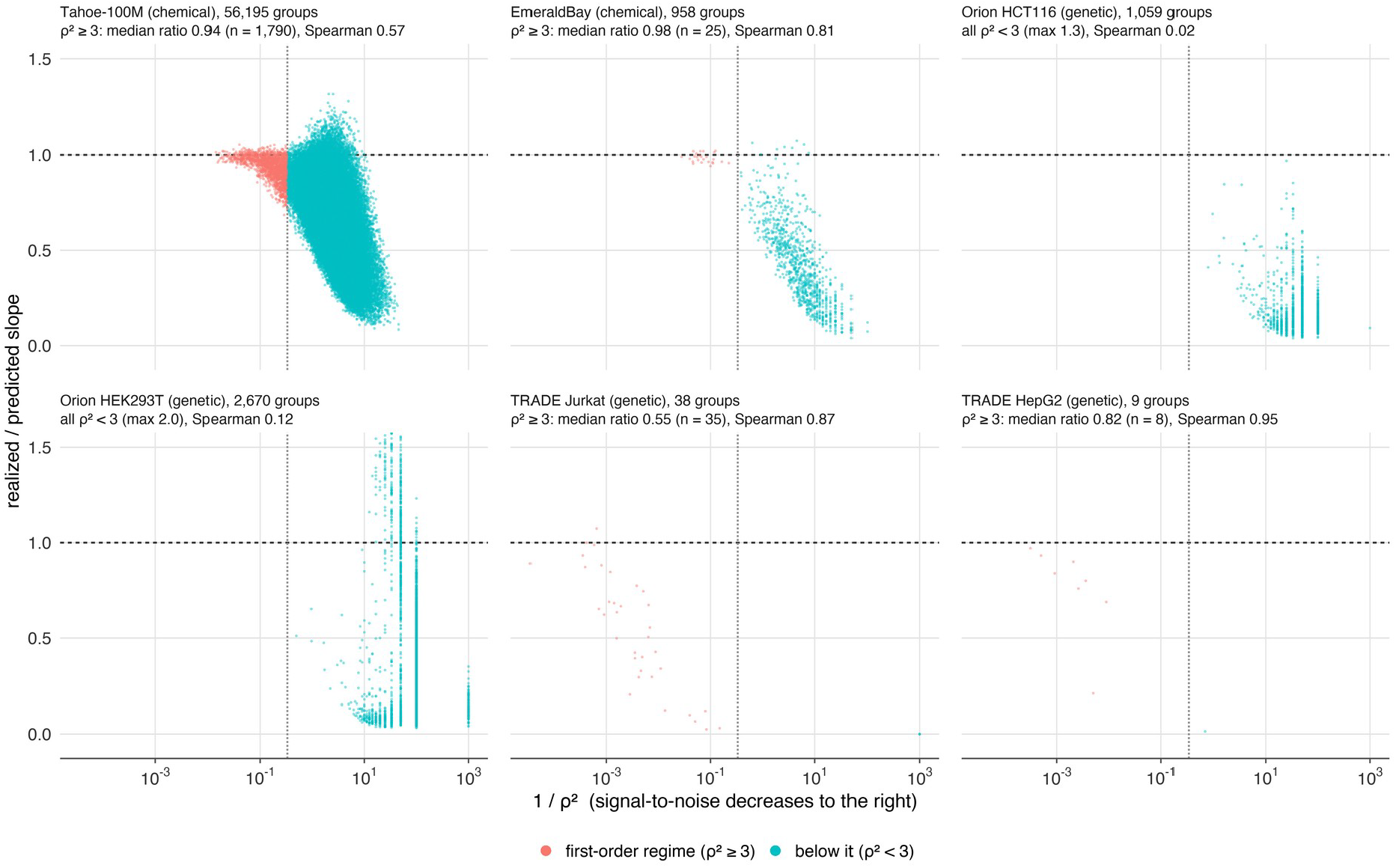
Held-out downsample-and-measure test on every group with *≥* 400 cells, one panel per screen. Each point is the realized/predicted slope ratio (no fitted parameter) against 1/ *ρ*^2^, so signal-to-noise decreases to the right. Colour separates the first-order regime (*ρ*^2^ *≥*3) from the regime below it. The dashed line marks a ratio of one and the dotted line *ρ*^2^=3. The ratio sits at one in the high-signal groups and falls monotonically as 1/ *ρ*^2^ grows.

## Discussion

This study makes explicit a quantity that direction-based perturbation screens have used only implicitly: the number of cells a condition needs before its perturbation direction can be trusted to a stated angular precision. We provide a single closed-form quota that converts a tolerance, a platform noise covariance, a condition’s effect magnitude and the control-pool size into a required treated depth, or into the conclusion that no treated depth suffices. Applied to six screens from four atlases, it shows that most contemporary data fall short. At a 5.7° tolerance the great majority of perturbations in today’s largest screens cannot resolve their direction, and the binding reason differs by design. On the genetic screens it is effect size and treated depth. On the largest chemical atlas it is a shared vehicle pool too small for the number of conditions it serves.

Sample sufficiency for direction depends on the match between each perturbation’s magnitude, the cells spent on it and the control it is referenced to, and it does not follow from the size of the atlas. Because *n*^*⋆*^ *∝* 1/ *m*^2^, the distribution of required cells follows from the effect-size distribution through an inverse-square law, and effect-size distributions are heavy at small values. The TRADE study quantifies this on the same genetic screens. A typical genome-wide knockdown changes about 45 genes whereas a typical essential-gene knockdown changes more than 500, and most of a perturbation’s transcriptome-wide impact resides in genes that never reach significance [36]. Our magnitude measures the same impact geometrically, so a magnitude distribution concentrated near zero forces a quota distribution far above realistic depth. The genome-wide and essential-gene screens fail the quota for opposite reasons, small *m* at a moderate depth of 155–204 cells versus large *m* at a very shallow 48–85 cells, and a screen’s regime is set the moment its library and its depth are chosen. Because raw displacement is inflated by sampling noise at shallow depth, as TRADE’s GATA1/EIF4A3 example shows in gene space [36], the quota subtracts the sampling-floor term from the squared magnitude on every dataset before it is applied.

The most portable quantity was *σ* ^2^, which stayed near 1.0 across a chemical platform at two timepoints and a genome-wide genetic platform. It is the residual cell-to-cell scatter after each condition’s mean is removed, in a fixed log-normalized, highly-variable-gene, 50-component embedding, so it reflects the measurement and not any particular perturbation. Where it moved, to 1.5–1.9 on the essential-gene screens, the cause was identifiable. Its value is set in part by the normalization scale of that embedding and must be re-estimated for each platform and pipeline. We measured the anisotropy directly. The within-condition covariance spans 3.6–5-fold across components on the chemical atlases, yet *tr* (*PΣ P*) varies by only 1.5–3% across conditions because drug displacements do not concentrate on the high-variance axes.

The quota complements existing design tools. scPower [23] and bulk replicate planning [24] target detection power, the multivariate analogue being a Hotelling test [37] whose non-centrality *v*^*⊤*^ *Σ*^*−*1^ *v* rewards low-noise directions. Directional precision is governed by a different functional, *tr* (*PΣ P*), which is increased by perpendicular noise, so a screen can be well powered to detect a shift yet under-powered to orient it. Neither tool sees the control arm. The two-arm quota, in contrast, exposes a small shared control as its own constraint through *m*_*min*_, which on the largest atlas dominates the result. The directional quota can serve as the sample-size standard for direction-based single-cell screens in the way replicate-number guidance did for bulk RNA-seq. It is closed-form, per-condition and computed from quantities a pilot already yields. Its payoff depends on the regime. Where the control is large, quota-guided allocation cuts the treated budget 2.8-fold (Table 3). On the pool-limited Tahoe atlas, adaptive and flat allocation recovered the same similarity graph and neither recovered mechanism labels, so there the quota serves to decide which conditions and which neighbour edges can be trusted and does not reallocate cells usefully.

Several limitations bound these conclusions. First, the noise model is the pooled within-condition covariance scaled by 1/ *n*. Between-well biological variation, plate effects and cell-cycle composition do not shrink with cell number and are not included, so *n*^*⋆*^ understates the requirement where they dominate, and multiplicative or rotational batch effects need dedicated integration upstream of the calculator [38,39]. Second, the downsample-and-measure test resamples the same cells that define its reference, so it verifies the finite-population sampling law and the small-angle linearization and does not test the independence of replicate wells. The over-sampled decision test likewise shares the control centroid between subsample and reference, so it does not exercise the control-pool term, and a replicate-well test of *m*_*min*_ remains to be done. Third, the quota is a first-order result whose guarantee holds for *ρ*^2^ well above one. Below the detection floor the calculator returns no count at all, and for the weak but detectable conditions the count it returns should be read as an order of magnitude and not as a precise requirement. Fourth, the tolerance *θ*_*⋆*_ =0.1 rad is a convention, and the sufficiency fractions move with it (Table 2). The criterion that matters for a given analysis is that each direction be resolved more tightly than the inter-mechanism angles it must separate. Fifth, the chemical spectra use a diagonal *Σ*, and the EmeraldBay and Tahoe headlines use different references (a per-line mean and a matched vehicle), so their rows in Table 1 are not directly comparable. Against its own two vehicle samples EmeraldBay would likely be largely pool-limited too. Sixth, the quota certifies the pseudobulk centroid direction only, and the compute savings from downsampling are polynomial. Finally, the machine-checked component is the deterministic algebra. The expectation and tail bound rest on symbolic and Monte-Carlo verification.

Three extensions follow. The per-cell guide abundance in FiCS Perturb-seq gives a continuous within-condition dose along which the 1/ *m*^2^ law could be tested within a single gene. A detection-mode companion using the Hotelling functional would report both whether an effect exists and how well its direction is resolved. A replicate-well calibration of the between-well variance would turn the within-condition quota into a mixed-model requirement.

## Conclusion

A single closed-form quota converts a tolerance, a platform noise covariance, a condition’s effect magnitude and the size of its control pool into the treated depth that resolves a perturbation direction, or into the finding that no treated depth does. Its deterministic core is machine-checked and its probabilistic content is verified by simulation. Applied to six screens from four atlases reprocessed through one embedding, the rule shows that most conditions in today’s largest screens cannot resolve their direction at a 5.7-degree tolerance. On the largest chemical atlas the binding constraint is the shared vehicle pool, and on the genome-wide genetic screens it is effect size and depth. The calculator gives experimenters this answer per condition before sequencing, and on the largest atlas it identifies the control pool, not the treated arm, as the first thing to enlarge.

## Supplemental Material

Supplemental Notes S1–S3 (the first- and second-order Delta expansion, the confidence quota and the marginal-variance cross-check), Supplemental Methods and Supplemental Tables S1–S4 are available online.

## Data and Code Availability

Tahoe-100M, EmeraldBay, X-Atlas/Orion and the TRADE essential-gene screens (GEO GSE264667) are publicly available datasets. The derived per-condition calibration fixtures used here, including the Orion and TRADE quota spectra, are included in the code repository. The calculator, the symbolic and Monte-Carlo verification suite, the Lean 4/Mathlib development, the from-raw recompute scripts and the manuscript-generation pipeline are openly available at https://github.com/huulocmedvnu/sample_sufficiency_calculator.

## Acknowledgments

This work received no specific funding, and the authors declare no competing interests. Anthropic’s Claude (used through the Claude Code environment) was employed under the authors’ direction to draft and revise the text of all sections of this article, to write and refactor the analysis, verification and figure code, and to check the reference list against the Crossref database. The study design, the data processing choices, the numerical results, the figures and the conclusions were specified, verified and approved by the authors, who take full responsibility for the content. Every text or citation edit proposed by the tool was reviewed against the primary sources before it was accepted. The tool was not used to generate or alter data or images, and it is not an author.

## Author Biographies

**Tran Thai Huu Loc** received the M.D. degree from the School of Medicine, Vietnam National University Ho Chi Minh City, Viet Nam, in 2019. After a year as a single-cell bioinformatician at BioTuring, San Diego, CA, USA (2019 to 2020), Dr. Tran practised obstetrics and gynecology at My Duc Hospital, Ho Chi Minh City (2021 to 2025), and since 2025 has been an obstetrician and gynecologist at the Department of Obstetrics and Gynecology, Hong Hung Hospital, Tay Ninh City, Viet Nam. Since 2026 Dr. Tran has also developed bioinformatics analyses for single-cell perturbation drug research with DataXight, Mountain View, CA, USA. Research interests include applications of artificial intelligence in obstetrics and gynecology and single-cell analysis. Dr. Tran has co-authored publications in Bioinformatics, Physiotherapy and Reproductive Sciences.

**Nguyen Thi Van** received the B.A. degree from the Banking Academy of Vietnam, Hanoi, Viet Nam, and the M.S. degree from the Department of Artificial Intelligence Convergence, Chonnam National University, Gwangju, Republic of Korea, in 2023, where she was a research assistant with the BIEL Laboratory, Chonnam National University Hwasun Hospital, working on deep learning for tumor detection in CT, MRI and PET imaging and on optical coherence tomography. She has since worked as a machine learning engineer on credit-risk models, as a research assistant with AI VIET NAM on semi-supervised segmentation of histopathology images, and as a bioinformatics research intern developing Nextflow pipelines for genomic diversity monitoring. Research interests include applied mathematics and statistics, bioinformatics, deep learning for medical imaging, and large language models in biomedical research.

