## Supplemental Material (Notes S1-S3, Methods, Tables S1-S4) for "How many cells resolve a perturbation direction? A closed-form, control-aware cell quota for single-cell perturbation screens"

### Supplemental Methods

#### Symbols

| symbol | meaning |
|---|---|
| $x_i^t, x_j^c$ | one treated / control cell, a $d$-vector in the embedding |
| $\hat{\mu}_t, \hat{\mu}_c$ | treated / control centroid (pseudobulk mean vector) |
| $v = \mu_t - \mu_c, \hat{v}$ | true / estimated perturbation vector |
| $m = \lVert v \rVert$ | effect magnitude |
| $u = v / m$ | unit direction (the mechanistic axis) |
| $\theta, \theta_{\cdot}$ | angular error; target root-mean-square tolerance |
| $\Sigma$ | per-cell within-condition covariance ($d \times d$) |
| $\sigma^2$ | scalar proxy: mean per-coordinate variance, $tr\,\Sigma / d$ |
| $S = \Sigma_t / n_t + \Sigma_c / n_c$ | sampling covariance of $\hat{v}$ |
| $P = I - u u^T$ | tangent-space projector, removes the noise along $u$ |
| $\rho^2 = m^2 / (u^T \Sigma u)$ | along-signal signal-to-noise ratio (per cell) |
| $d_{\text{eff}}$ | effective number of contributing noise dimensions |
| $N_0$ | cells already acquired for a condition |
| $n_c$ | control-pool size |
| $n_t^{\cdot}$ | required treated cells (the quota) |
| $m_{min}$ | control-pool floor, $\sqrt{tr(P \Sigma P) / (n_c \theta_{\cdot}^2)}$ |

#### Pipeline topology

The order of operations matters. (i) The PCA basis is fitted once on the pooled single-cell expression matrix of each atlas, defining a shared 50-dimensional coordinate space. (ii) The per-cell covariance $\Sigma$ is estimated from individual embedded cells within that space as the pooled within-condition residual covariance: each condition is centred on its own mean and the residuals are pooled with $\sum(n-1)$ degrees of freedom. (iii) Only afterwards is each condition summarized by the coordinate mean of its embedded cells. Aggregating counts before PCA would leave a single point per condition and collapse $\Sigma$ to zero, removing the functional $tr(P\Sigma P)$ on which the quota depends. Because $n_t \neq n_c$ in general, the two cell matrices are not subtracted directly. Each is collapsed to a centroid vector first.

The chemical pipelines (Tahoe-100M, EmeraldBay) read per-condition sufficient statistics (sums, sums of squares and counts in the 50-dimensional space) and therefore carry a diagonal $\Sigma = diag(\ell_1, \ldots, \ell_{50})$ of per-component within-condition variances. The genetic pipelines (Orion, TRADE) carry the full $50 \times 50$ matrix. On the chemical atlases the per-component variance spans 3.6–5-fold, yet $tr(P\Sigma P)$ varies by only $\pm 1.5$–3% across conditions (Tahoe 45.8–47.0 against $tr\,\Sigma = 47.8$), because drug displacement directions do not concentrate on the high-variance axes. On Orion $tr(P\Sigma P)$ ranges 30–45.

For Tahoe-100M the vehicle centroid for each (plate × line) pools every DMSO_TF well on that plate (two wells on 11 plates, three on 3). A condition spanning several plates is referenced to the treated-weighted average of its plate vehicles, with the variance-equivalent pool size $n_c = \left(\sum_p c_p\right)^2 / \sum_p c_p^2 / n_{c,p}$. The EmeraldBay embedding uses a frozen 2,000-gene highly variable set, because per-fit selection made the 50-dimensional subspace unstable under resampling (mean $\cos^2 = 0.79$ between disjoint-shard refits, against 0.96 with the frozen set).

#### Plug-in estimation

In practice $\Sigma$ and $u$ are estimated from a pilot. The plug-in quota $\hat{n}^*$ uses the pooled within-condition covariance $\hat{\Sigma}$ (in the PCA basis, its diagonal) and the pilot direction $\hat{u} = \hat{v}/\|\hat{v}\|$, with $tr(P\hat{\Sigma}P) = \sum_k \left(1 - \hat{u}_k^2\right)\ell_k$. It is consistent, with leading error $O\left(\sqrt{d/n_{\text{pilot}}}\right)$ from the covariance and $O(\theta_{\text{pilot}})$ from the direction. The trace is first-order sensitive to direction error, $\delta\, tr(P\Sigma P) = -2\delta u^T \Sigma u + O(\|\delta u\|^2)$, which vanishes only when $u$ is an eigenvector of $\Sigma$, so pilot direction error propagates into the quota at first order as run-to-run scatter. The bias is second order because $\hat{u}$ is unbiased to first order. Ledoit–Wolf shrinkage [1] can regularize $\hat{\Sigma}$ when $d$ is not small relative to the pilot.

**Within-condition versus marginal variance**

The derivation calls for the within-condition residual covariance, the cell-to-cell scatter left after removing each condition's own mean, because that is the noise that enters a centroid. The within-condition per-cell variance on Tahoe-100M is $\sigma^2 = 0.9567$ over all 95,624,334 cells. The coarser marginal variance of all cells in the embedding, which additionally absorbs between-condition and between-line structure, is 2.406 and yields a quota about 2.5-fold larger. The within-condition Tahoe value nearly coincides with EmeraldBay's (0.9174), a match across timepoints on the same platform to within about 4%. On both Orion lines the marginal variance nearly equals the within-condition value (0.917 versus 0.913, and 1.040 versus 1.033).

**Verification and continuous integration**

The formula is checked four independent ways, all exercised by an automated test suite (26 tests: unit and property tests, golden-value tests that lock the within-condition $\sigma^2$ of every dataset and the headline spectrum, an integration test on held-out EmeraldBay groups, and two guards that fail if the quota is re-implemented outside the engine) alongside a Lean 4/Mathlib development, both run in continuous integration on Python 3.10–3.12.

- **Symbolic Jacobian.** A SymPy-derived Jacobian of the normalization map matches the analytic $1/mP$ exactly. The residual is the zero matrix, with maximal floating-point deviation $2.8 \times 10^{-17}$.

- **Monte-Carlo consistency.** With $K=50{,}000$ samples ($d=50$, $d_{\text{eff}}=12.2$, $\rho^2=3.82\times10^5$) the relative error of $E[\theta^2]=tr(PSP)/m^2$ is 0.13%. A second harness at moderate signal-to-noise pins the second-order expansion of Supplemental Note S1 to below 0.5%, against about 3% for the lever-arm-only form.
- **Laurent–Massart tail.** The empirical exceedance is $Pr(\theta>\theta_{\star})=0.34\%\leq\delta=10\%$.
- **Held-out integration test.** On a locked set of high-cell-count EmeraldBay groups the predicted curve $\theta^2(n)=tr(P\Sigma P)/m^2(1/n-1/N)$ reproduces the observed downsampling with mean relative error 0.6–5.3%, spanning the strong-signal fit ( $R^2>0.997$) and a low-magnitude group in the breakdown regime ($R^2=0.979$).
- **Formal proof.** The deterministic core, the Jacobian $D(x\mapsto x/\|x\|)|_v=1/\|v\|P$, the trace identity $tr(P\Sigma P)=tr\,\Sigma-u^T\Sigma u$, its isotropic reduction $\sigma^2(d-1)$ and its diagonal form $\sum_k(1-u_k^2)\ell_k$, is machine-checked in Lean 4 against Mathlib (pinned to v4.31.0) with no `sorry` and only the standard axioms. The probabilistic results (the expectation and the tail bound) are not formalized.

The embedding was built with scanpy 1.12.2 (seurat-flavour highly variable genes, arpack PCA) on Python 3.13.12 with numpy 2.4.4, scipy 1.18.0, scikit-learn 1.9.0 and anndata 0.12.18. The environment is pinned in the repository. The manuscript is typeset with pandoc 3.10 and tectonic 0.15.

##### Held-out protocol details

For each group the reference direction uses all $N$ acquired cells relative to the per-line mean (a large, near-noiseless reference distinct from the plate-matched vehicle used for the spectrum). Depths follow a geometric grid capped at $N/2$ so that every fitted point stays away from the degenerate boundary $n=N$, at which the subsample is the whole pool and

$\theta \equiv 0$. Tahoe-100M uses $R = 120$ subsamples per depth and the other atlases $R = 300$. The angle is the arccosine of the cosine similarity between the subsample and reference unit vectors, clipped to $[-1, 1]$. The realized statistic is the root-mean-square angle over subsamples, and the fit is ordinary least squares of $\theta^2$ on $(1/n - 1/N)$ through the origin. Because subsamples are drawn without replacement from the cells that define the reference, this test verifies the finite-population sampling law and the small-angle linearization. It does not test the independence of replicate wells.

The over-sampled decision test downsamples each condition predicted OVER to $n = \lceil n^{\star} \rceil$ (200 subsamples) and requires the realized root-mean-square angle against the full-depth direction to be within $1.05\,\theta_{\star}$. On EmeraldBay it was run at $\theta_{\star} = 0.20$ rad over the 3,971 (condition × line) groups with at least 100 cells (706 predicted OVER, 706 met). On Tahoe-100M it was run under the per-line-mean reference over conditions with at least 400 cells (13,964 of 13,964), and against the pooled shared vehicle over all 6,467 predicted-OVER conditions (6,442 met, with a median realized angle of 0.060 rad and a maximum of 0.113 rad). The 25 exceptions have median bias-corrected $m = 4.4$ and $n^{\star} \approx 240$. Of these, 23 fall in NCI-H460 and 2 in hTERT-HPNE.

#### Supplemental Note S1: First-order Delta expansion and its second-order correction

Only noise perpendicular to the signal rotates the direction, and the first-order rule makes this precise. Because the Jacobian of the normalization map $x \mapsto x/\|x\|$ at $x = v$ is exactly $1/m P$, only the perpendicular part of the noise reaches $\hat{u}$. Here $P = I - u u^T$ is the projector onto the tangent space of the unit sphere at $u$ (rank $d - 1$, $Pu = 0$). To first order $\hat{u} - u = 1/m P e$, which is orthogonal to $u$, so $\theta \approx \|Pe\|/m$ and $E[\theta^2] = tr(PSP)/m^2$.

The exact geometry makes the regime of validity explicit. Decomposing the sampling error $e = \hat{v} - v$ into its along- and across-signal parts, $e = (u^T e) u + P e$, the estimated vector is $\hat{v} = (m + u^T e) u + P e$, so the exact angular error is

$$\tan\theta = \frac{\|P e\|}{m + u^T e}.$$

The denominator is a fluctuating lever arm: the across-signal noise fixes the transverse displacement, while the along-signal noise lengthens or shortens the effective magnitude against which it is measured. To first order this dependence is inert, because $P$ annihilates the along-signal component exactly. Freezing the lever arm at $m$ recovers $\theta \approx \|P e\|/m$, asymptotically exact when $\rho^2 \equiv m^2/(u^T S u) \gg 1$, where the relative lever-arm fluctuation has standard deviation $1/\rho \ll 1$. Writing $r \equiv \|P e\|/m$ and $s \equiv u^T e/m$, the expansion $\tan\theta = r/(1+s)$ with $\theta^2 = \tan^2\theta - 2/3 \tan^4\theta + O(\rho^{-6})$ gives, after taking expectations with Isserlis' theorem,

$$E[\theta^2] = \frac{tr(P S P)}{m^2}\left(1 + \frac{1}{m^2}\left[3 u^T S u + \frac{6 u^T S P S u}{tr(P S P)} - 2/3\, tr(P S P) - \frac{4\, tr\left((P S P)^2\right)}{3\, tr(P S P)}\right] + O(\rho^{-4})\right).$$

The first two bracket terms are the lever-arm correction (positive, $O(\rho^{-2})$ relative to the leading term): a shortened lever amplifies a fixed transverse displacement more than an equally lengthened one attenuates it. The last two are the arc-versus-tangent correction (negative, $O(\theta_\star^2)$ relative): the arc is shorter than the tangent. Both must be kept, because retaining only the lever-arm group over-predicts the angle at this order. At the quota the arc-versus-tangent group dominates, so the relative correction is $O(\theta_\star^2)$, below 1% at $\theta_\star = 0.1$ rad, and its net sign is a shortfall of the realized angle below the first-order prediction. Monte-Carlo pins the corrected expression to below 0.5%. This second-order structure is what produces the low signal-to-noise slope deficit of Figure 5. The exact $\theta^2(n)$ is gently concave in $(1/n - 1/N)$, so a straight-line fit returns a slope biased below the small-noise limit by an amount that grows as the subsampling reaches lower depth, while remaining linear through the origin at high $R^2$. Outside the small-angle regime ($\rho \lesssim 1$) the angle is near-uniform on the sphere and no finite quota clears the noise. The correct report is then “not resolvable at this depth”, and no number should be given.

#### Supplemental Note S2: Distribution of the angular error and the confidence quota

Because $\|Pe\|^2$ is a quadratic form in a Gaussian, it is a generalized chi-square, $\|Pe\|^2 \overset{d}{=} \sum_{i=1}^{d-1} v_i z_i^2$ with $z_i \overset{\text{iid}}{\sim} N(0,1)$ and $v_i$ the eigenvalues of $PSP$ [2]. Hence

$$\theta^2 \approx \frac{1}{m^2} \sum_{i=1}^{d-1} v_i z_i^2, E[\theta^2] = \frac{tr(PSP)}{m^2}, Var(\theta^2) = \frac{2\,tr\left((PSP)^2\right)}{m^4}.$$

The effective degrees of freedom $d_{\text{eff}} = (tr\,PSP)^2 / tr\left((PSP)^2\right) \le d-1$ count how many noise dimensions effectively contribute [3]. For the single-cell data studied here it is much smaller than $d-1$ (12.2 in the Monte-Carlo check) because a few axes dominate the variance.

To guarantee $Pr(\theta > \theta_{\cdot}) \le \delta$ we apply the Laurent–Massart tail bound for weighted chi-square statistics, the specialization of the Hanson–Wright inequality to a weighted sum of chi-squares [4–6]:

$$Pr\left(\|Pe\|^2 \ge tr(PSP) + 2\|PSP\|_F \sqrt{L} + 2\|PSP\|_{op} L\right) \le e^{-L}, L = \log(1/\delta).$$

Requiring $m^2\theta_{\cdot}^2$ to exceed the right-hand side and inverting for $n$ with $S = 2\Sigma/n$ gives the closed-form confidence quota

$$n_\delta^{\cdot} = \frac{2}{m^2\theta_{\cdot}^2}\left[tr(P\Sigma P) + 2\|P\Sigma P\|_F \sqrt{\log 1/\delta} + 2\|P\Sigma P\|_{op} \log 1/\delta\right] = n_{\text{aniso}}^{\cdot}\left(1 + O\left(\sqrt{\log(1/\delta)/d_{\text{eff}}}\right)\right),$$

the mean quota inflated by a multiplicative safety margin set by $d_{\mathrm{eff}}$ and the confidence level, not a different scaling law. Empirically the margin is conservative: at $\delta = 10\%$ the realized exceedance is 0.34%.

One qualification applies. The inequality rigorously controls the perpendicular-noise energy $\|Pe\|^2$ and becomes a statement about the angle only through the first-order identification $\theta \approx \|Pe\|/m$. As Note S1 shows, the true angle also depends on the along-signal fluctuation in the denominator, which the bound does not account for. The confidence quota is therefore an approximate $(1-\delta)$ guarantee on $\theta$, exact as $\rho \to \infty$, not a certified finite-sample bound on the geometric angle. In practice the Laurent–Massart constants are loose enough to dominate the neglected term, so the error is in the safe direction for planning.

#### Supplemental Note S3: Marginal-variance cross-check

The main text uses the within-condition residual variance $\sigma^2 = 0.9567$. As a conservative check one may instead use the marginal per-cell variance $\sigma^2 = 2.406$, which additionally absorbs between-condition and between-line structure and over-states the noise that enters a centroid. Because both the quota and the control-pool floor scale with $\sigma$ ($m_{min} \propto \sigma$), substituting the marginal value raises every floor by $\sqrt{2.406/0.9567} \approx 1.6$-fold and can only increase the control-pool-limited fraction. The qualitative conclusion, that most conditions are unresolvable against Tahoe's shared vehicle at a stringent tolerance, is unchanged and if anything strengthened.

#### Supplemental tables

**Supplemental Table S1. Datasets, controls and calibration constants** ($\theta_{\cdot} = 0.1$ rad). $\sigma^2$ is the within-condition per-cell variance, C is the large-control bound $(d-1)\sigma^2/\theta_{\cdot}^2$, and $m_{min}$ is the median control-pool floor at the real $n_c$. The regime column is over / under / pool-limited over all conditions for Tahoe and over / under among detectable knockdowns for the genetic screens.

| dataset | platform, modality | cells (post-QC) | conditions | control ($n_c$) | $\sigma^2$ | C | $m_{min}$ | median $m$ | median depth | detectable % | regime |
|---|---|---|---|---|---|---|---|---|---|---|---|
| Tahoe-100M | Vevo Mosaic, chemical (24 h) | 95,624,334 | 56,827 | pooled plate DMSO_TF (median 3,113) | 0.9567 | 4,688 | 1.22 | 1.096 | 1,296 | 94.4 | 11.4 / 25.2 / 55.4 |
| EmeraldBay | Vevo Mosaic, chemical (5 d) | 1,831,648 | 4,912 | per-line mean (about 22,800) | 0.9174 | 4,495 | 0.45 | 1.175 | 240 | 87.6 | 2.7 / 82.5 / 0.5 |
| Orion HCT116 | Xaira FiCS, genome-wide CRISPRi | about 4 million | 17,585 | NTC pool (165,562) | 0.9133 | 4,475 | 0.16 | 0.141 | 155 | 14.7 | 0.0 / 100.0 |
| Orion HEK293T | Xaira FiCS, genome-wide CRISPRi | about 4 million | 17,856 | NTC pool (218,838) | 1.0334 | 5,063 | 0.12 | 0.351 | 204 | 35.3 | 0.1 / 99.9 |
| TRADE Jurkat | essential-gene CRISPRi | about 250,000 | 2,187 | NTC pool (11,514) | 1.4975 | 7,338 | 0.78 | 1.841 | 85 | 70.6 | 1.2 / 98.8 |
| TRADE HepG2 | essential-gene CRISPRi | about 250,000 | 1,883 | NTC pool (4,380) | 1.8667 | 9,147 | 1.39 | 2.249 | 48 | 60.9 | 4.4 / 95.6 |

**Supplemental Table S2. Per-drug sampling-sufficiency spectrum across all doses and cell lines** ($\theta_*=0.1$ rad, Tahoe-100M against its pooled shared DMSO vehicle, counts out of the 150 conditions per drug = 3 doses × 50 lines). Because the vehicle floor $m_{min}=1.22$ exceeds the median effect magnitude, POOL-LIMITED is the dominant column: 240 of 379 drugs are pool-limited in a majority of their conditions, 9 are over-sampled in a majority, and 23 are over-sampled in no condition. Median *m* is bias-corrected, and median $n^*$ is over the resolvable conditions. Selected rows shown.

| Drug (class) | median $m$ | median $n^*$ | OVER | UNDER | Ghost | POOL-LIM | not-det |
|---|---|---|---|---|---|---|---|
| Panobinostat (HDAC) | 3.87 | 294 | 121 | 10 | 0 | 15 | 4 |
| Homoharringtonine (protein synthesis) | 5.64 | 125 | 111 | 20 | 0 | 18 | 1 |
| Harringtonine (protein synthesis) | 3.44 | 250 | 82 | 23 | 1 | 38 | 6 |
| Idarubicin (TOP2A) | 2.70 | 348 | 77 | 29 | 1 | 37 | 6 |
| Trametinib (MEK) | 2.11 | 1,148 | 93 | 35 | 1 | 18 | 3 |
| Palbociclib (CDK4/6) | 1.34 | 4,680 | 8 | 68 | 1 | 67 | 6 |
| 4EGI-1 (eIF4E) | 1.30 | 5,474 | 7 | 63 | 2 | 72 | 6 |
| (S)-Crizotinib (ALK/MET) | 0.94 | 8,885 | 1 | 18 | 5 | 115 | 11 |

**Supplemental Table S3. Sampling summary by dose** (all 379 drugs × 50 lines at each dose, Tahoe-100M, pooled shared DMSO vehicle). The five regime percentages plus not-detectable sum to 100. Median *m* is bias-corrected (raw per-dose medians 1.11 / 1.13 / 1.22).

| dose | conditions | % OVER | % UNDER | % Ghost | % POOL-LIM | % not-det | median $m$ |
|---|---|---|---|---|---|---|---|
| 0.05 µM | 18,948 | 6.9 | 24.0 | 2.5 | 60.8 | 5.8 | 1.061 |
| 0.50 µM | 18,929 | 13.7 | 26.9 | 2.4 | 51.1 | 5.9 | 1.071 |
| 5.00 µM | 18,950 | 13.5 | 24.8 | 2.2 | 54.2 | 5.3 | 1.171 |

**Supplemental Table S4. Exemplars for the downsampling-invariance test** (EmeraldBay, $\theta_{*}=0.1$ rad, two-arm quota with the real per-line control pool, bias-corrected $m$). Downsampling both groups to 92 cells leaves the OVER centroid stable (centroid–baseline distance changed by $\leq 0.6\%$, nearest-neighbour cosines by $\leq 0.005$, replicate SD 0.008) but destabilizes the UNDER one (distance +19%, from 0.78 to 0.93, cosines $-0.10$ to $-0.12$, replicate SD 0.10).

| condition (× cell line) | $N_0$ | $m$ | $n^{*}$ (cells) | diagnostic |
|---|---|---|---|---|
| DMSO_T0 × HS-578T (reference population) | 1,067 | 5.89 | 92 | OVER (11.6× spare) |
| Gemcitabine × MIA PaCa-2 | 280 | 0.69 | 42,666 | UNDER (152× short) |
